# Deletion of PIEZO1 in adult cardiomyocytes accelerates cardiac aging and causes premature death

**DOI:** 10.1101/2025.01.20.633994

**Authors:** Ze-Yan Yu, Yang Guo, Scott Kesteven, Delfine Cheng, Hanzhou Lei, Jianxin Wu, Evelyn Nadar, Peter Macdonald, Munira Xaymardan, Charles D. Cox, Michael P. Feneley, Boris Martinac

## Abstract

Mechanosensitive PIEZO1 channels have emerged as key transducers of mechanical forces in the cardiovascular system. In cardiomyocytes, we previously showed that PIEZO1 decodes mechanical cues driving pressure-overload induced hypertrophy. However, conflicting reports exist on the influence of PIEZO1 on baseline cardiac function. Here we show that conditional deletion of *Piezo1* from cardiomyocytes in adult mice results in premature mortality. The hearts from these mice exhibited signs of accelerated aging, including elevated markers of the senescence associated secretory phenotype, with significant blunting of the normal cardiac hypertrophic response to aging, associated with a reduction in the activation of the pro-hypertrophic Ca^2+^/calmodulin-dependent protein kinase II (CaMKII). Functionally, aged-*Piezo1* KO mice exhibited impaired cardiac relaxation due to altered cellular Ca^2+^ handling kinetics. Young adult *Piezo1* KO mice exhibited a normal resting heart rate but developed significant progressive sinus bradycardia and cardiac fibrotic remodelling with aging, which was most prominent in the right atrium, where *Piezo1* expression is highest in the healthy heart. Mechanistically, loss of PIEZO1 was associated with a marked reduction in the anti-fibrotic molecule, atrial natriuretic peptide (ANP). Moreover, *in vivo,* ANP release instigated by atrial stretch was markedly blunted in conditional *Piezo1* KO mice, providing a plausible and long sought-after mechanism for the link between mechanical stretch and ANP release. Taken together, our data show that PIEZO1 is a crucial homeostatic molecule during cardiac aging, enabling adaptation to an aging tissue microenvironment.

## Introduction

The cardiovascular system provides a pulsatile blood supply throughout the body. Concurrently, this pulsatile action applies dynamic mechanical forces on cells and tissues within the cardiovascular system. Cells adapt to these ever-changing mechanical signals by sensing and responding to these mechanical cues using molecular sensors of mechanical forces ^1^. Since the discovery of mechanosensitive PIEZO channels in 2010^2^, there has been growing evidence for the role of PIEZO1 in sensing mechanical cues relevant to both cardiovascular physiology and pathology^1,3–8^.

PIEZO1 channels are expressed in almost all cell types within the cardiovascular system, including throughout the myocardium in endothelial cells (ECs), cardiac fibroblasts (CFs) and cardiomyocytes (CMs)^5,9–12^. Early studies indicated that PIEZO1 expression was low in adult murine heart tissue and even lower in adult cardiomyocytes^9,11,12^. Despite this fact, it is now clear that PIEZO1 is indeed present in the sarcolemma of adult cardiomyocytes, where it plays a crucial role in mechanotransduction^11–13^.

Initial studies suggested *Piezo1* was dispensable for normal cardiac development^4,5,11^ but cardiomyocyte-specific conditional deletion of *Piezo1* during development led to enlargement of the heart and the development of a dilated cardiomyopathy-like phenotype^11^. Conversely, genetically engineered overexpression of PIEZO1 in cardiomyocytes resulted in severe cardiac hypertrophy and arrhythmias, which worsened with age^11^. Consistent with this hypertrophic phenotype with PIEZO1 overexpression, we have previously demonstrated that left ventricular hypertrophy (LVH) in response to pressure overload induced by transverse aortic constriction (TAC) was severely blunted by conditional deletion of cardiomyocyte *Piezo1* in the adult heart^12^. The downstream activation of LVH by PIEZO1 was dependent on a histone deacetylase 4 (HDAC4), myocyte enhancer factor 2 (MEF2), and Ca2+/calmodulin-dependent kinase II (CaMKII) signalling pathway. These results showed that PIEZO1 is the mechanosensor which converts the elevated myocardial forces caused by pressure overload into chemical signals that trigger hypertrophic signalling cascades, which in turn produce pathological LVH^12,13^.

Despite the established role of PIEZO1 as a key transducer of mechanical forces in the cardiovascular system^1^, conflicting reports clearly exist regarding how cardiomyocyte expressed PIEZO1 channels affect baseline cardiac function. To explore this, we deleted *Piezo1* conditionally in the cardiomyocytes of adult mice and tracked their cardiac function as they aged. Here we demonstrate that conditional deletion of *Piezo1* in adult mouse cardiomyocytes resulted in premature mortality. Hearts from these mice exhibited signs of accelerated aging, manifested as rises in markers of the senescence associated secretory phenotype, with blunting of the normal age-related cardiac hypertrophic response due to reduced activation of the pro-hypertrophic kinase CaMKII, and impaired cardiac relaxation related to abnormalities in cellular Ca^2+^ handling. *Piezo1* KO mice also displayed an age-induced bradycardia and cardiac fibrosis that was most prominent in the right atrium, where *Piezo1* expression is highest in the healthy heart. Mechanistically, the expression of the anti-fibrotic protein atrial natriuretic peptide (ANP) was significantly reduced in *Piezo1* KO mice and its release in response to *in vivo* volume-overload, which generates acute atrial stretch, was also markedly reduced in *Piezo1* KO mice. These findings unveil PIEZO1 as an important homeostatic molecule in cardiac aging and provide a plausible connection between PIEZO1 and the molecular mechanisms enabling cardiac adaptation to the aging tissue microenvironment.

## Materials and Methods

### Mice

All experimental procedures were approved by the Animal Ethics Committee of Garvan/St Vincent’s (Australia), in accordance with the guidelines of both the Australian code for the care and use of animals for scientific purposes (8th edition, National Health and Medical Research Council, AU, 2013) and the Guide for the Care and Use of Laboratory Animals (8th edition, National Research Council, USA, 2011). Male mice were utilised throughout, and all were on the C57BL/6J genetic background.

As previously described^12^, to produce *Piezo1^flox/flox^; αMHC-MCM^+/-^* (termed *P1^fl/fl^MCM^+/-^*) mice, the inducible cardiomyocyte-specific Piezo1 knockout (KO) mice were generated by crossing homozygous *Piezo1*^flox/flox^ mice (The Jackson Laboratory Stock, No: 029213) with homozygous *Myh6*-MerCreMer mice (MCM), which have a tamoxifen-inducible Cre recombinase under the control of the α-myosin heavy-chain (αMHC; *Myh6)* promoter^14^. The age and sex-matched *Piezo1^wt/wt^; αMHC-MCM^+/-^* (termed *αMHC-MCM^+/-^*) mice were used as controls for experiments. For six consecutive days, male *P1^fl/fl^MCM^+/-^*mice aged 8–10 weeks were given a daily intraperitoneal (*I.P*.) injection of tamoxifen (30 mg/kg, Sigma, T5648) dissolved in 95% peanut oil (Sigma, P2144) to induce Cre recombinase-mediated deletion of exons 20–23 of the *Piezo1* gene selectively in adult mouse cardiomyocytes (referred to as Piezo1 KO). The age- and sex-matched *P1^wt/wt^αMHC-MCM^+/-^*mice treated with tamoxifen served as Cre-only controls. *P1^fl/fl^MCM^+/-^*mice treated with peanut oil acted as flox controls. Mice were given 10 days to recover after the last injection of tamoxifen before experiments.

### Echocardiographic measurements

As previously described^15^, echocardiography was performed using an MS400 18–38 MHz transducer probe and VEVO 2,100 ultrasound system (VisualSonics Inc., Canada). The mice were anesthetized (3–5% isoflurane for induction, 1–2% isoflurane for maintenance with adjustment to maintain heart rate at ∼500 bpm) and imaged at 3-, 6-, 12-, and 18 months after *Piezo1* KO to assess cardiac function.

To investigate the influence of adrenergic drive on heart rate 18 months after tamoxifen treatment, the heart rates of *αMHC-MCM^+/-^* and *Piezo1* KO mice were measured by echocardiography before (baseline) and then 20 minutes after intraperitoneal injection of 1.0 mg/kg of Metoprolol (Sigma, USA), a selective β-adrenoceptor blocker. The acquisition of images and evaluation of data were performed by an operator blinded to treatment.

### Mouse heart rate measurements

We employed the CODA tail-cuff system (Kent Scientific, Torrington, CT, USA) to assess the heart rate of conscious mice. Mice were acclimated to the restraint tubes and tail-cuff system daily for three consecutive days before measurements to minimize stress-induced variability. This acclimatization involved placing each mouse in the restraining tube for 15 minutes and attaching the tail cuff without inflating it to familiarize the animals with the setup. On the day of measurement, each mouse was placed in a restraint tube, and the tail was secured in the CODA tail-cuff sensor. The system was set to measure the heart rate using a non-invasive volume pressure recording technique that detects blood flow in the tail^16^. Measurements were performed in a quiet room at a controlled temperature of 25 °C to prevent temperature-induced changes in cardiovascular parameters. The procedure included three preliminary inflation-deflation cycles to stabilize measurements, followed by 10 cycles to obtain consistent heart rate readings. The mean heart rate was calculated from the 10 cycles after excluding outliers and artifacts. All heart rate data were recorded and analyzed using the CODA software. To ensure accuracy and consistency, we measured the heart rate in each mouse three times on three separate days and use the average as its heart rate data. For comparison, additional heart rate measurements were taken from mice under anaesthesia with isoflurane, allowing for assessment of heart rate under both conscious and unconscious states.

### Invasive hemodynamic measurements

As previously described^17,18^, mice were anesthetized by inhalation of isoflurane (1.5%) and a 1.4F micro-tip pressure catheter (Millar Instruments Inc, Houston, Texas, USA) was inserted into the left ventricle via the right carotid artery. The heart rate, aortic systolic pressure, LV systolic pressure, +dP/dt, and –dP/dt were recorded (LabChart 6 Reader, AD Instruments, P/L). Animals were sacrificed, and the heart weight (HW), left ventricular weight (LVW), right ventricular weight (RVW), left atrial weight (LAW), and right atrial weight (RAW), normalized to body weight (BW) and to tibia length (TL) were measured as indicators of hypertrophy.

### Mouse LV cardiomyocyte isolation and purification

As previously described^18^, the mice were heparinized and euthanized according to the Animal Research Act 1985 No 123 (New South Wales, Australia). Hearts were dissected and perfused through the aorta and the coronary arteries with 10 mL pH 7.2 perfusion buffer containing 135 mM NaCl, 4 mM KCl, 1 mM MgCl_2_, 0.33 mM NaH_2_PO_2_, 10 mM HEPES, 10 mM Glucose, 10 mM 2,3-Butanedione 2-monoxime (BDM), and 5 mM Taurine, with a Langendorff apparatus at 37 °C for 5 minutes. Next, 30 mL digestion buffer composed of the above solution and Collagenase B, D (dose by BW: 0.4 mg/g, Roche) and Protease Enzyme Type XIV (dose by BW: 0.07 mg/g, Sigma-Aldrich) was used to perfuse the hearts for 15 minutes. After the perfusion, the heart was removed from the setup and placed into a pH 7.4 transfer buffer (Buffer A) containing 135 mM NaCl, 4 mM KCl, 1 mM MgCl_2_, 0.33 mM NaH_2_PO_2_, 10 mM HEPES, 5.5 mM Glucose, 10 mM BDM, and 5 mg/mL BSA. Both atria and the right ventricle were discarded (other than for Figure 8k where each individual chamber was separated to isolate individual cardiomyocytes), and the LV muscle was torn into small pieces and gently triturated in transfer buffer with a pipette to isolate cardiomyocytes. The suspension was then filtered through a 200 micro falcon cup filter (BD) and centrifuged at 20xg for 2 minutes. After that, the cardiomyocytes were used either for Ca^2+^ imaging experiments or for purification described previously^19,20^, which confirmed that rod-shaped cardiomyocytes accounted for more than 85% of the total purified cardiomyocytes. The isolated cardiomyocytes were frozen immediately in liquid nitrogen and stored at -80 °C for following experiments.

### Cardiomyocyte Ca^2+^ transient measurements

For the Ca^2+^ imaging experiments, a transfer buffer (Buffer B) at pH 7.4 was prepared, containing 135 mM NaCl, 5.4 mM KCl, 0.5 mM MgCl2, 1.8 mM CaCl2, 10 mM HEPES, 5.5 mM glucose, 10 mM BDM, and 5 mg/mL BSA. Buffer A and B were mixed in various ratios to create solutions with Ca^2+^ concentrations of 0.06, 0.24, 0.6, and 0.8 mM. To prevent intracellular Ca^2+^ overload, isolated mouse ventricular myocytes were gradually transferred through these solutions to adapt to a final Ca^2+^ concentration of 0.8 mM. The cells were then plated into a 96-well microplate (Greiner Bio-One, Germany) at a density of 1500 cells per well and incubated with 2.5 µM of a fluorescent Ca^2+^ indicator, Cal-520 AM (Abcam, ab171868, UK) at 37°C for 40 minutes. After incubation, the cells were rinsed and maintained in 100 µL of a Buffer A and B mixture containing 0.8 mM Ca^2+^. Ca^2+^ imaging was conducted using a Nikon Eclipse Ti2-E Inverted epifluorescence microscope (Nikon Instruments, Japan) at a frame rate of 100 frames per second. Data recording lasted for a total of 30 seconds, during which electrical pacing of 60 V and 0.5 Hz was applied 10 times starting from the fifth second using a bipolar stimulator (manufacturer to be added later). Using a method similar to that described by Guo et al.,^10^ Ca^2+^ transient data was initially analyzed with the built-in NIS-Elements Microscope Imaging Software, version 5.11.03 (Nikon Instruments, Japan), to extract the intensity profiles over time. Following this, custom MATLAB (MathWorks, USA) software, developed by Anton Shpak and Satya Arjunan at the Innovation Centre, Victor Chang Cardiac Research Institute, Australia, was utilized to measure Ca^2+^ transients and calculate parameters such as Ca^2+^ transient duration (CTD), rise time, fall time, and others. The software also corrected baseline intensity rundown caused by sample bleaching during microscopy.

### Atomic force micrscopy

Freshly isolated mouse ventricular myocytes were plated on coated FluoroDishes (FD35-100, WPI) coated with Geltrex (Gibco, A1413201, USA) 1:100 in serum free M199 media (Sigma-Aldrich, M4530, USA), for 1 h at 37 °C. Excess Geltrex was aspirated, and cells were plated and left to adhere for ∼1.5 h, in transfer buffer (Buffer A), before stiffness measurement. Cell stiffness was measured using an atomic force microscope (JPK Nanowizard 4, Bruker) at 37°C and a 5 mm spherical probe (SAA-SPH-5um, Bruker). After calibration of the probe, the following settings were used: setpoint of 0.5 nN, z length of 4 µm and z speed of 0.5 µm/s. Force curves were acquired from the middle of cells and analysed using the Hertz/Sneddon model with a Poisson’s ratio of 0.4. We analysed between 43-53 individual cardiomyocytes per heart and plotted the average value for 3 biological replicates per condition.

### Quantitative Reverse Polymerase Chain Reaction (qRT-PCR)

Gene expression was determined by quantitative reverse qRT-PCR. Total RNA was extracted and purified from LV tissue and isolated cardiomyocytes with the RNeasy Fibrous Tissue Mini Kit (QIAGEN), following the manufacturer’s protocol. RNA (500 ng) was reverse transcribed into cDNA using the SuperScript III First-Strand Synthesis SuperMix kit (Invitrogen). cDNA was subjected to PCR amplification to detect *Piezo1*, *ANP* (*Nppa*), *collagen III* (*Col3α3)*, *SERCA2a, phospholamban (PLN), Slc8a1, RyR2, p16, p53, p21, interleukin (IL)1β, IL6, transforming growth factor β (TGFβ), tumour necrosis factor α (TNFα)* gene expression, using the PCR master mix LightCycler 480 SYBR Green I Master (Invitrogen) and performed with the CFX384 Touch Real-Time PCR Detection System (Bio-Rad). The mRNA expression levels were normalized to those of GAPDH to calculate relative gene expression using the delta-delta Ct method. The mouse RT-PCR primers (Sigma-Aldrich) used are shown in Supplementary Table 1.

### Western blotting

For total protein extraction, LV tissue and isolated cardiomyocytes were lysed in a pH 7.4 lysis buffer containing 150 mM NaCl, 50 mM Tris-HCL, 1% Triton X-100, 1 mM sodium orthovanadate, 1 mM beta-glycerophosphate, 5 mM dithiothreitol and MiniComplete protease inhibitors (Roche). Briefly, LV tissue was lysed using NE-PER nuclear and cytoplasmic extraction reagents (Pierce Biotechnology) and Protease Inhibitor Cocktail Kit and Halt Phosphatase Inhibitor Cocktail (Pierce Biotechnology), both with a homogenizer (PRO Scientific). Total protein (30 μg for each sample) was loaded on 4%-20% Mini-PROTEAN TGX Gels (Bio-Rad) and separated by electrophoresis. Samples were transferred to PVDF membranes (Bio-Rad), blocked with 5% bovine serum albumin (BSA) then labelled overnight with primary antibodies except for anti-SERCA2a for 1 hour at 4 °C (Extended Data Table 4): anti-SERCA2a (1:35000, Abcam), anti-Phospholamban (1:1500, Abcam), anti-Phospho-Phospholamban (Thr17, 1:1500, Badrilla), anti-NCX1 (1:1000, Thermo Scientific), anti-CaMKIIδ (1:1000; Abcam), anti-p-CaMKII (Thr287, 1:5000; Thermo Scientific), Anti-GAPDH (1:10000; Cell Signaling Technology) was used to standardize sample loading. Horseradish peroxidase-conjugated (HRP) goat anti-rabbit (1:10000) secondary antibodies (Abcam) (Extended Data Table 4) were used at room temperature for one hour. Immunologic detection was accomplished using SuperSignal™ West Pico PLUS

Chemiluminescent Substrate (Thermo Scientific). Protein levels were quantified by densitometry using ImageJ (NIH) software. Protein levels were normalized to relative changes in GAPDH and expressed as fold changes relative to those of control animals.

### Histological and immunofluorescence analysis

As previously described^12,18^, Masson’s trichrome stain was used to quantify fibrosis (collagen fibres stain blue). The hearts were excised from isoflurane-euthanized mice and washed with phosphate-buffered saline (PBS). Hearts were then cut longitudinally in the coronal plane, embedded into optimal cutting temperature (OCT) compound (Sakura Finetek), and gradually frozen in melting isopentate, precooled in liquid nitrogen to avoid tissue damage, and stored at -80 °C for following experiments. Serial 6 μm sections were sliced with a cryostat (Leica) and stained using a Masson’s trichrome staining kit (Sigma-Aldrich), following the manufacturer’s instructions. Images of the LV, RV, LA or RA were obtained with 4 to 6 fields per section^21^ using a brightfield microscope (Leica). Blue-stained areas of fibrosis within sections were determined using colour-based thresholding^22^ and measured with ImageJ software (NIH; http://rsbweb.nih.gov/ij/). The percentage of total fibrotic area was calculated by taking the sum of the blue-stained areas divided by the total LV, RV, LA and RA area.

### Acute expansion of intravascular volume

Three-month-old male *αMHC-MCM^+/-^* mice and *Piezo1 KO* mice received whole blood from wild-type donor mice via a right jugular vein catheter similar to methods previously described^23^. Briefly, the donor mice were anesthetized with 5% isoflurane and ventilated at 120 breaths per min (Harvard Apparatus Rodent Ventilator), and the blood was drawn by cardiac puncture using a heparinized syringe. The recipient mice were canulated using a jugular catheter inserted into right jugular vein under isoflurane anaesthesia delivered through intra-tracheal intubation. 1.5 ml heparinized whole blood was infused into mice over 3 minutes, which expanded the mouse blood volume by an estimated 60%. 150 µl of blood was collected prior to infusion and 5 and 15 minutes after infusion by cardiac puncture into chilled haematocrit tubes containing 5 µl of 0.2 M sodium-EDTA and was immediately centrifuged at 4 °C. Plasma was removed and stored at –80 °C ready to assay for ANP levels.

### Plasma ANP assay

Plasma ANP levels were measured with Mouse Atrial Natriuretic Peptide ELISA kit (Cusabio), following the manufacturer’s instructions. Analysis of plasma ANP level was performed by means of radioimmunoassay methods (Peninsula Laboratories, Belmont, CA, USA). Values of ANP were expressed as pg/mL.

### Statistics

All experiments and analyses were blinded. Averaged data are presented as mean ± standard error of the mean (SEM). The statistical analyses were performed using GraphPad Prism software, version 7.04 (GraphPad). For comparisons among three or more sets of data with one factor or two factors, one-way or two-way ANOVA was used accordingly, followed by Tukey’s post-hoc test. For comparisons between two groups, Welch’s T-test, two-tailed was used, and *p* < 0.05 was considered statistically significant. All samples used in this study were biological repeats, not technical repeats. All experiments were conducted using at least two independent materials to reproduce similar results.

## Results

### Cardiomyocyte-specific deletion of Piezo1 in adult mice causes premature death

We utilized previously established conditional *Piezo1* knockout (*Piezo1* KO) mice^12^ to achieve targeted deletion of *Piezo1* in adult cardiomyocytes through tamoxifen induction (**Fig. 1a**). This model was compared with two age-matched control groups: *P1^fl/fl^MCM^+/-^*mice treated with oil and *αMHC-MCM^+/-^* mice treated with tamoxifen, as detailed in the **Materials and Methods**. Our findings demonstrated that *Piezo1* KO mice began to experience premature death from around 19 months of age, 72 weeks after tamoxifen treatment. Survival rates declined to 65.8% by 22 months of age, 78 weeks after tamoxifen induction. This was significantly lower than the survival rates observed in the *P1^fl/fl^MCM^+/-^* (90.6%, p = 0.01) and *αMHC-MCM^+/-^* (93.8%, p = 0.004) control groups (**Fig. 1b**). These results suggest that cardiomyocyte PIEZO1 may play an important protective role during cardiac aging.

**Figure. 1.**
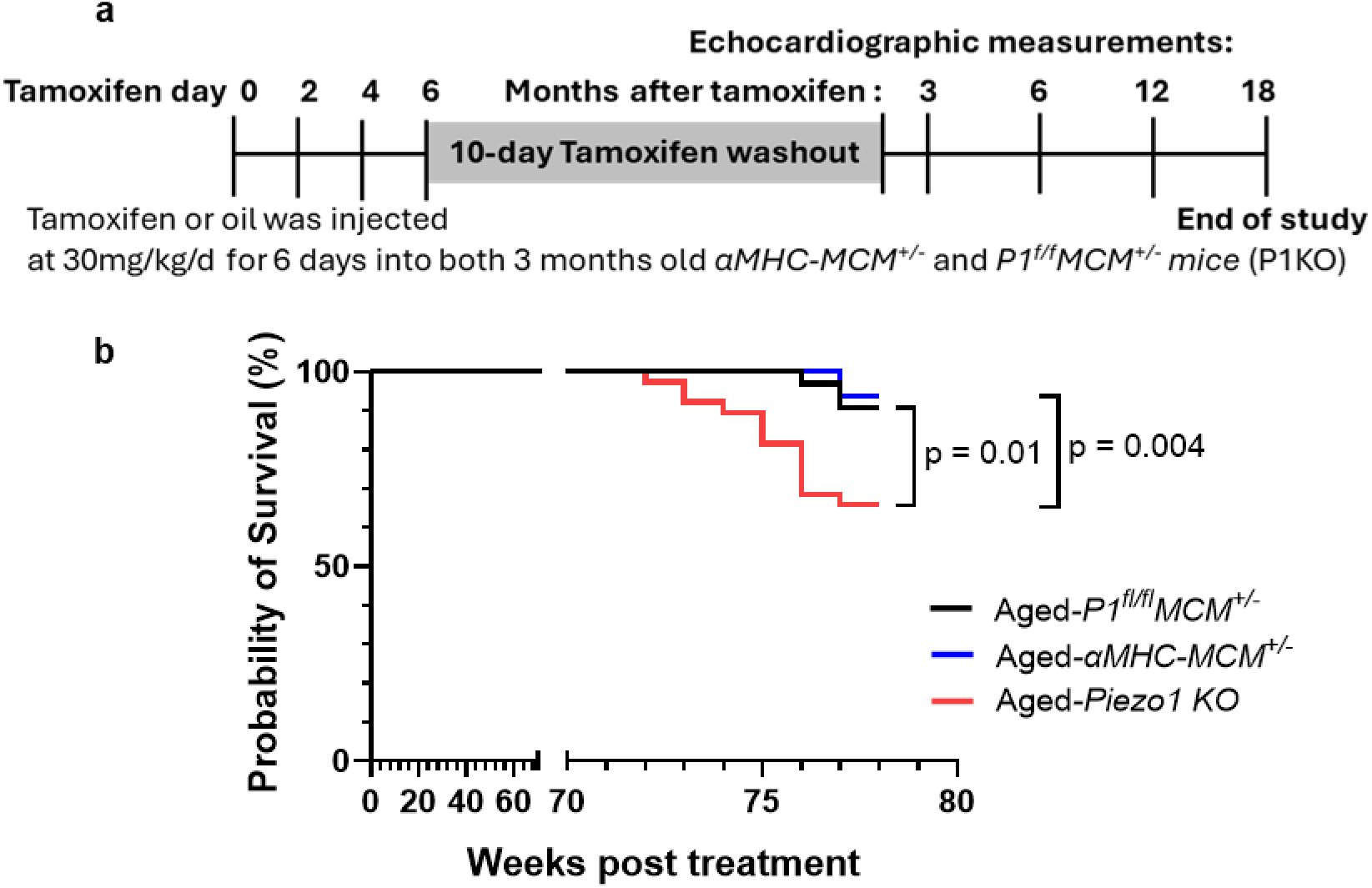
Deletion of *Piezo1* in cardiomyocytes results in premature death in aging mice. (**a**) Experimental design and timeline for tamoxifen induction and serial echocardiographic measurements. (**b**) Kaplan-Meier curve illustrating increased mortality in 21-month-old *Piezo1* KO (n=13 out of 38) mice (Aged-*Piezo1 KO)* 18 months after deletion of *Piezo1* compared to Aged-*αMHC-MCM^+/-^* (n=2 out of 32) and Aged-*P1^fl/fl^MCM^+/-^ (n=3 out of 32)* control groups.

### Cardiomyocyte-specific deletion of Piezo1 in adult mice causes accelerated cardiac senescence

Accumulating evidence suggests that cellular senescence plays a pivotal role in various heart diseases^24,25^. Cardiac aging is recognized as a primary driver of morbidity and mortality^26–28^. We hypothesized that the early mortality observed in aging *Piezo1* KO mice may be associated with premature cardiac senescence. To test this, we first examined the expression levels of the three classic senescence markers, *p16*, *p53* and its downstream target, *p21,* in left ventricular (LV) tissue of aged (21 months old) *αMHC-MCM^+/-^* control hearts. These were compared to both healthy young (3 months old) *αMHC-MCM^+/-^* control hearts and age-matched *Piezo1* KO hearts.

Quantitative reverse transcript (qRT)-PCR revealed that aged *Piezo1* KO hearts had considerably higher mRNA expression of p16, p53, and p21 than age-matched *αMHC-MCM+/-* control hearts (**Fig. 2a-c**). In contrast, *p16* expression was only increased in aged control hearts as opposed to young control hearts (**Fig. 2a**), suggesting that *Piezo1* deletion in cardiomyocytes exacerbated senescence. To further confirm our findings, we assessed the expression levels of senescence associated secretory phenotype (SASP) factors, such as pro-inflammatory cytokines interleukin-1 (I*L1β*), *IL6*, transforming growth factor-*β* (TGF*β*), and tumour necrosis factor-α (TNFα), which are known to cause inflammation and contribute to age-related cardiovascular disease^24,29^. mRNA levels of *IL1β, IL6, TGFβ and TNFα* levels were significantly elevated in aged *αMHC-MCM^+/-^* control hearts in comparison to young *αMHC-MCM^+/-^*control hearts (**Fig. 2d-g**), but aged *Piezo1* KO hearts showed significantly higher levels of expression than age-matched control hearts (**Fig. 2d-g**). These results imply that PIEZO1 has a protective function and that its loss in cardiomyocytes accelerates aging related processes.

**Figure. 2.**
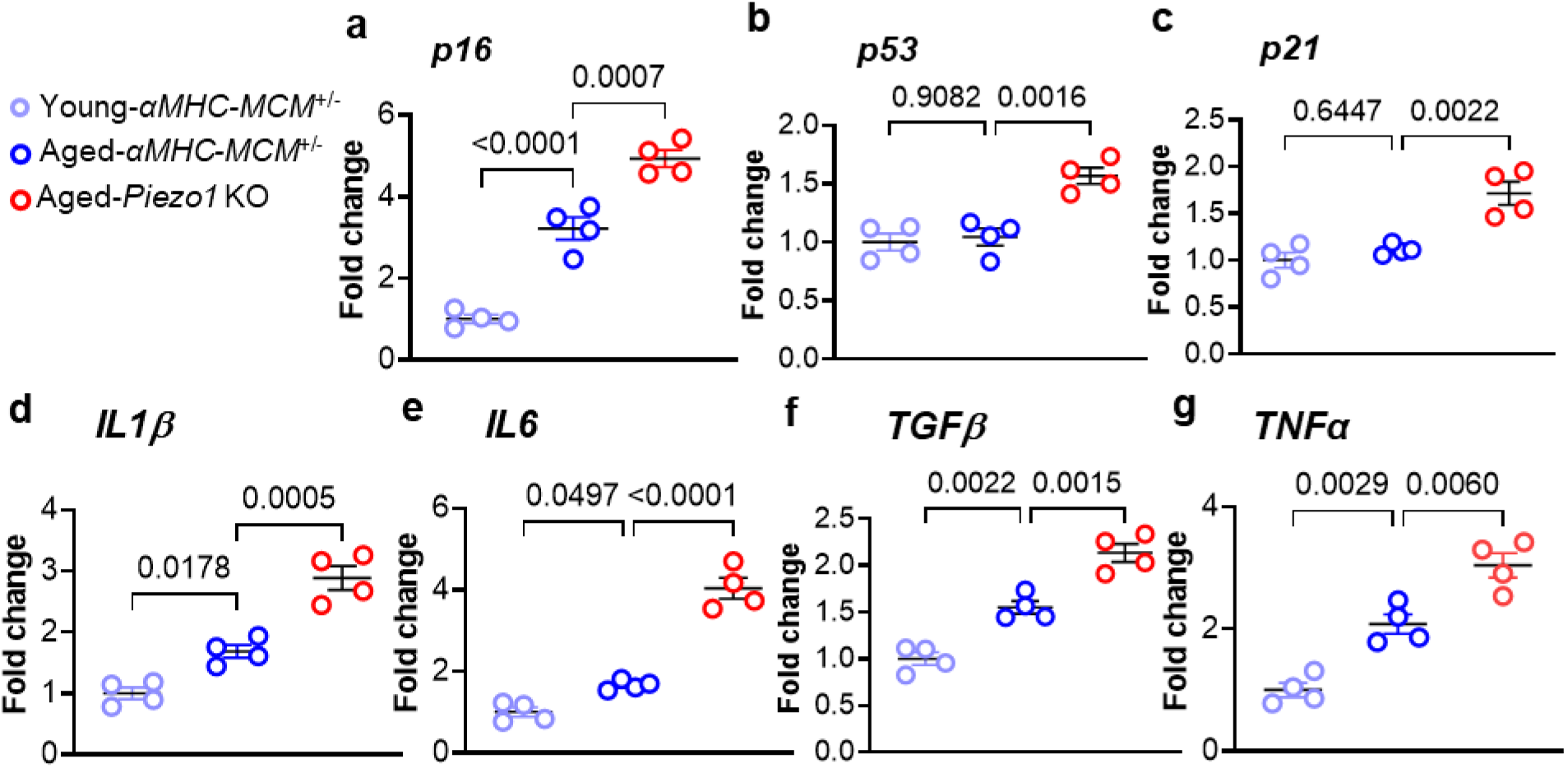
Deletion of *Piezo1* in adult cardiomyocytes results in an increase in senescence associated secretory phenotype (SASP) factors indicative of accelerated aging. Relative mRNA expression of senescent genes measured in 3-month-old tamoxifen treated *αMHC-MCM^+/-^* mice (Young *αMHC-MCM^+/-^* mice) and 21-month-old mice 18 months following tamoxifen treatment in both *αMHC-MCM^+/-^* mice (Aged-*αMHC-MCM^+/-^*) and Aged-*Piezo1* KO mice. (**a**) *p16*, (**b**) *p53*, (**c**) *p21*, (**d**) *IL1β*, (**e**) *IL6*, (**f**) *TGFβ*, (**g**) *TNFα*. The mRNA relative expression was normalized with GAPDH and calculated as fold change relative to Young-*αMHC-MCM^+/-^* mice (n = 4 per group). Results are presented as mean ± SEM. One-way ANOVA with Tukey’s post-hoc test for multiple comparisons was used.

### Cardiomyocyte-specific deletion of Piezo1 in adult mice inhibits the normal hypertrophic response to aging

Cardiac aging is accompanied by cardiac hypertrophy^30^. We conducted post-mortem examinations to determine whether this behaviour is present in aged *Piezo1* KO mice. There were no significant differences in body weight between genotypes within the young or aged study groups (**Fig. 3a**). As expected, we observed significant hypertrophy in aged mice compared to young controls (**Fig. 3b-f**). This included changes in all cardiac chambers except for the right ventricle (**Fig. 3g**). Interestingly, we found that the age-related hypertrophy was significantly blunted in *Piezo1* KO mice hearts, which were approximately 12% lighter than those of both the age-matched *P1^fl/fl^MCM^+/-^* and *αMHC-MCM^+/-^*control groups (**Fig. 3b**). This observation was further supported by normalized ratios of heart weight to tibia length (HW/TL; ∼11-12 % smaller in *Piezo1* KO), left ventricle weight to tibia length (LVW/TL; ∼15-16 % smaller in *Piezo1* KO), left atrial weight to tibia length (LAW/TL; 23-27 % smaller in *Piezo1* KO), and right atrial weight to tibia length (RAW/TL; 22-24 % smaller in *Piezo1* KO) (**Fig. 3c-e**).

**Figure. 3.**
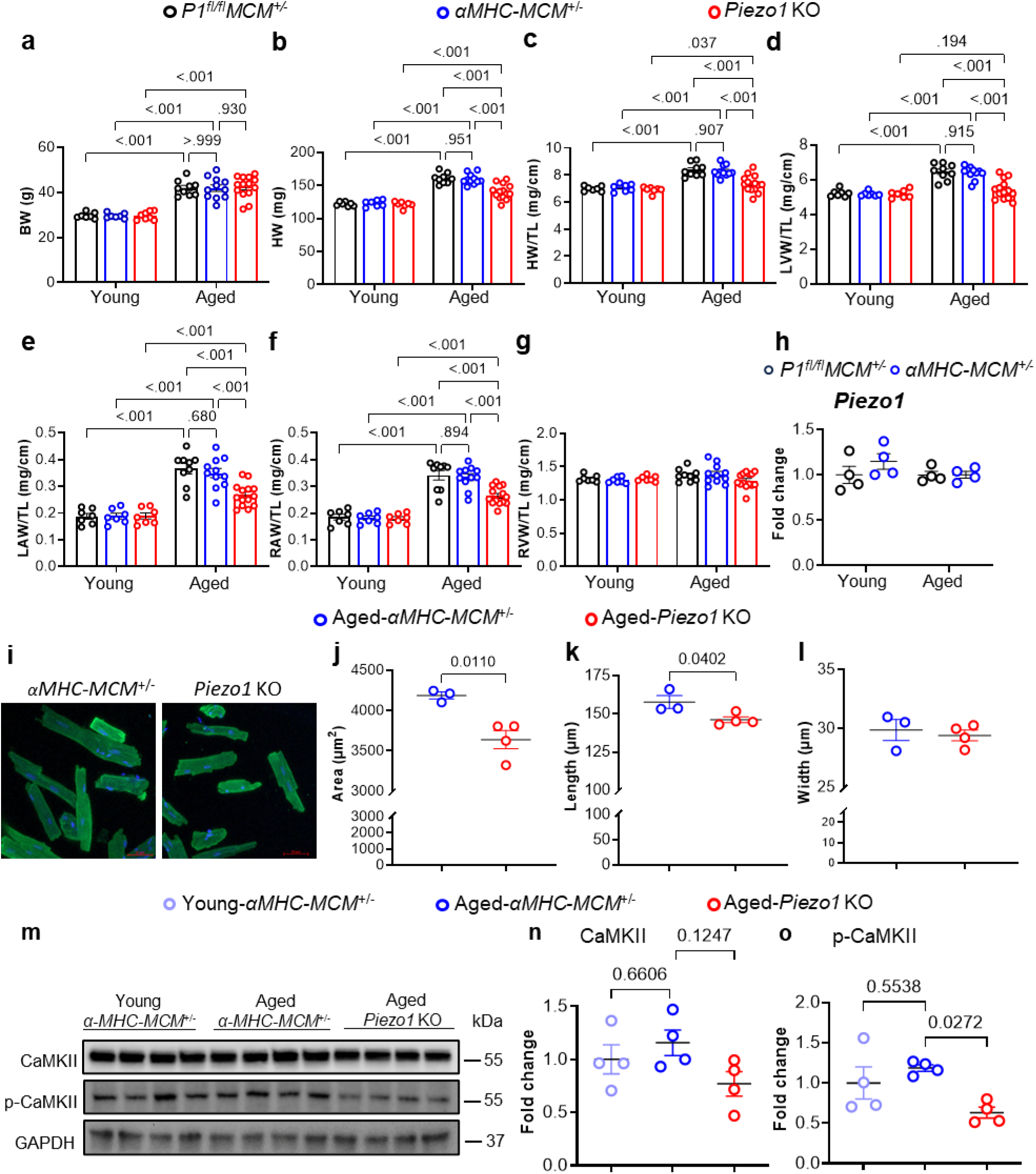
Deletion of *Piezo1* in adult cardiomyocytes inhibits the normal hypertrophic response to aging. Anatomical measurements from 3-month-old mice around 2 weeks following tamoxifen treatment (*Young*) and 21-month-old mice 18 months following tamoxifen treatment (*Aged*) in both *αMHC-MCM^+/-^* and *Piezo1* KO mice, and oil treated *P1^fl/fl^MCM^+/-^*controls showing; (**a**) BW, Body weight. (**b**) HW, heart weight. (**c**) HW/TL, heart weight to tibia length. (**d**) LVW/TL, left ventricular weight to tibia length. (**e**) LAW/TL, left atrial weight to tibia length. (**f**) RAW/TL, right atrial weight to tibia length. (**g**) RVW/TL, right ventricular weight to tibia length (n = 9 – 14 per group). (**h**) Relative *Piezo1* mRNA expression in LV tissues from 3-month-old and 21-month-old after 18 months following oil treated *P1^f/l/fl^MCM^+/-^*mice and tamoxifen treated *αMHC-MCM^+/-^* mice (n = 4 per group). The mRNA relative expression was normalized with GAPDH and calculated as fold change relative to 3-month-old oil treated *P1^fl/fl^MCM^+/-^* mice. (**i**) Representative immunofluorescence images of single cardiomyocytes (green, cardiac troponin T; blue, DAPI for nuclei) from 21-month-old mice 18 months after tamoxifen treatment in both *αMHC-MCM^+/-^*(n = 3) and *Piezo1* KO hearts (n = 4) (scale bar = 50 µm). (**j – l**) Quantitative analysis of cardiomyocytes (number of cells for the measurements ranging from 60 – 70 per heart) showing (**j**) surface area; (**k**) length; (**l**) width. (**m**) Representative western blots showing the expression of CaMKII and p-CaMKII in 3-month-old tamoxifen treated *αMHC-MCM^+/-^* mice (*Young*) and 21-month-old mice 18 months following tamoxifen treatment in both *αMHC-MCM^+/-^* (*Aged*), and *Piezo1 KO* mice (*Aged*). (**o - p**) Quantitative data of CaMKII (**o)** and p-CaMKII (**p)** were normalized with GAPDH. Fold changes were calculated relative to 3-month-old tamoxifen treated *αMHC-MCM^+/-^* mice (n = 4 per group). Results are presented as mean ± SEM, One-way ANOVA with Tukey’s post-hoc test for multiple comparisons was used.

To determine whether *Piezo1* gene expression levels were changed with age, we measured mRNA levels of *Piezo1* in the LV tissues of our two control groups (*P1^fl/fl^MCM^+/-^* or *αMHC-MCM^+/-^*) at 3- and 18-months following treatment either with oil or tamoxifen. We observed no significant changes in mRNA levels of *Piezo1* across the examined groups (**Fig. 3h**), illustrating myocardial *Piezo1* levels do not change significantly during aging. To further characterise the blunting of age-induced hypertrophy in *Piezo1* KO hearts, we isolated individual cardiomyocytes from the left ventricle. Freshly isolated cardiomyocytes from aged *Piezo1* KO mice hearts were approximately 13% smaller than age-matched *αMHC-MCM^+/-^* control hearts (**Fig. 3i-l**). This reduction in area was mediated by a significant reduction in cell length (**Fig. 3k**) rather than cell width (**Fig. 3l**). This observation was consistent with the smaller weights of *Piezo1* KO mice hearts.

We have previously shown that PIEZO1 mediated activation of Ca^2+^/calmodulin-dependent protein kinase II (CaMKII) drives cardiac hypertrophy in response to increased afterload^12,13,17,18^. We postulated that CaMKII activation might also drive the hypertrophy observed during aging and thus the absence of CaMKII activation in the absence of PIEZO1 might explain the blunting of age-related cardiac hypertrophy observed in aged *Piezo1* KO hearts. To verify this, we used western blots to quantify the amount of CaMKII in aged *αMHC-MCM^+/-^* control LV tissue compared with both young *αMHC-MCM^+/-^* LV tissue (3 months old) and aged *Piezo1* KO LV tissue. While there was no significant change in total CaMKII expression between the *αMHC-MCM^+/-^*controls and aged *Piezo1* KO groups (**Fig. 3m-n**), there was a significant reduction in activated CaMKII, which is auto-phosphorylated at threonine 287, in the aged *Piezo1* KO LV tissue when compared to the *αMHC-MCM^+/-^*group (**Fig. 3m-o**). This is congruent with a model where during aging, PIEZO1 activity drives CaMKII signalling resulting in hypertrophy. In the absence of PIEZO1 in our *Piezo1* KO mice, this signalling is absent and results in reduced hypertrophy with aging.

### Cardiomyocyte-specific deletion of Piezo1 in adult mice impairs cardiac relaxation during aging

To evaluate whether the deletion of cardiomyocyte PIEZO1 impacts cardiac function during aging, echocardiography was performed on *Piezo1* KO mice at 3, 6, 12 and 18 months after tamoxifen induction, in comparison to the age-matched control groups. Between 3 to 12 months following tamoxifen treatment, there were no significant changes in echocardiographic parameters across the study groups (Supplementary Fig. 1). Similarly, no changes in left ventricular ejection fraction (LVEF) were detected among the three age-matched groups 18 months after tamoxifen treatment (**Fig. 4a**). While there were no significant differences in cardiac output between aged *P1^fl/fl^MCM^+/-^*mice and *αMHC-MCM^+/-^* control mice, the *Piezo1* KO mice displayed a ∼30% lower cardiac output compared to *P1^fl/fl^MCM^+/-^*control mice **Fig. 4b**), but this resulted mainly from a significant reduction in heart rate (data to be presented in more detail in Fig.7).

**Figure. 4.**
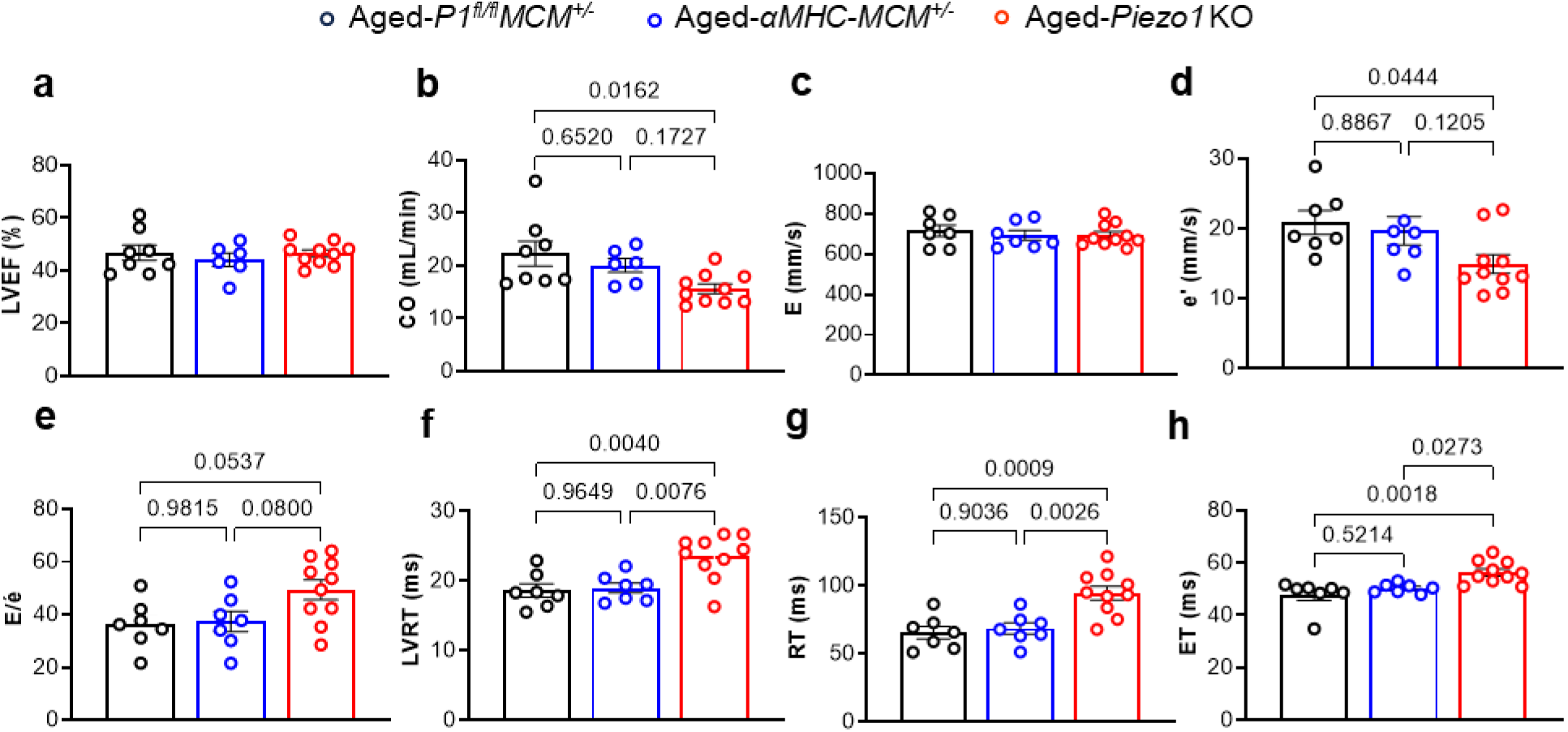
Deletion of *Piezo1* in adult cardiomyocytes impairs cardiac relaxation during aging. (**a**-**h**) Echocardiographic measurements in 21-month-old mice 18 months following tamoxifen treatment in both *αMHC-MCM^+/-^* and *Piezo1 KO* mice, and oil treated *P1^fl/fl^MCM^+/-^* controls showing; (**a**) LVEF, LV ejection fraction. (**b**) CO, cardiac output. (**c**) E, early diastolic peak transmitral flow velocity. (**d**) e’, early diastolic peak mitral annual velocity. (**e**) E/e’ ratio (**f**) IVRT, LV isovolumic relaxation time. (**g**) RT, LV relaxation time. (**h**) ET, LV ejection time (7 – 10 per group).

We next evaluated diastolic LV function by measuring E (early diastolic peak transmitral flow velocity), e’ (early diastolic peak mitral annulus velocity), and the E/e’ ratio, a key parameter for grading diastolic dysfunction^31^. Our results showed no statistical differences in the E values among the groups, but the e’ value in the *Piezo1* KO group was 28% lower than that in the *P1^fl/fl^MCM^+/-^* control mice, driving a 36% higher E/e’ ratio in *Piezo1* KO mice (**Fig. 4c-e**). *Piezo1* KO mice also exhibited a 26% longer left ventricular isovolumic relaxation time (IVRT), a 44% longer relaxation time (RT) and a 18% longer ejection time (ET) compared to *P1^fl/fl^MCM^+/-^*control mice. Similarly, *Piezo1* KO mice exhibited a 24% longer IVRT, a 37% longer RT and a 12% longer ET compared to *αMHC-MCM^+/-^* control mice (**Fig. 4f-h**). Together, these results demonstrate that deletion of cardiomyocyte *Piezo1* impairs cardiac relaxation in aged mice.

### Cardiomyocyte-specific deletion of Piezo1 in adult mice impairs Ca^2+^ handling kinetics in aged cardiomyocytes

To begin to understand the cellular basis of the effects of PIEZO1 on cardiac relaxation in aged mice, we first asked the question of whether the deletion of cardiomyocyte PIEZO1 alters Ca^2+^ handling in aged cardiomyocytes. We compared Ca^2+^ transients in freshly isolated ventricular myocytes from aged *Piezo1* KO hearts with the age-matched *αMHC-MCM^+/-^* control hearts, under 60 V electrical pacing at 0.5 Hz. Representative Ca^2+^ transients from one *αMHC-MCM^+/-^* control ventricular myocyte and one *Piezo1* KO ventricular myocyte are shown in **Fig. 5a**. Quantitative analyses revealed no significant difference in the Ca^2+^ transient duration at 30% recovery (CTD30) between the aged *Piezo1* KO cardiomyocytes and age-matched *αMHC-MCM^+/-^* control cardiomyocytes, but CTD50 was significantly prolonged by 29% and CTD90 was significantly prolonged by 42% in aged *Piezo1* KO cardiomyocytes, with a significant reduction in the CTD 50/90 ratio (**Fig. 5 b-e**). Additionally, while there was no significant difference in the rise time between 10% and 90% of the peak of the Ca^2+^ transient, the fall time between 90% and 10% from the peak was 47% longer in aged *Piezo1* KO cardiomyocytes (**Fig. 5f-g**). These findings indicate that the deletion of cardiomyocyte *Piezo1* causes abnormalities of Ca^2+^ handling kinetics in aged cardiomyocytes that are consistent with the observed impairment of cardiac relaxation.

**Figure. 5.**
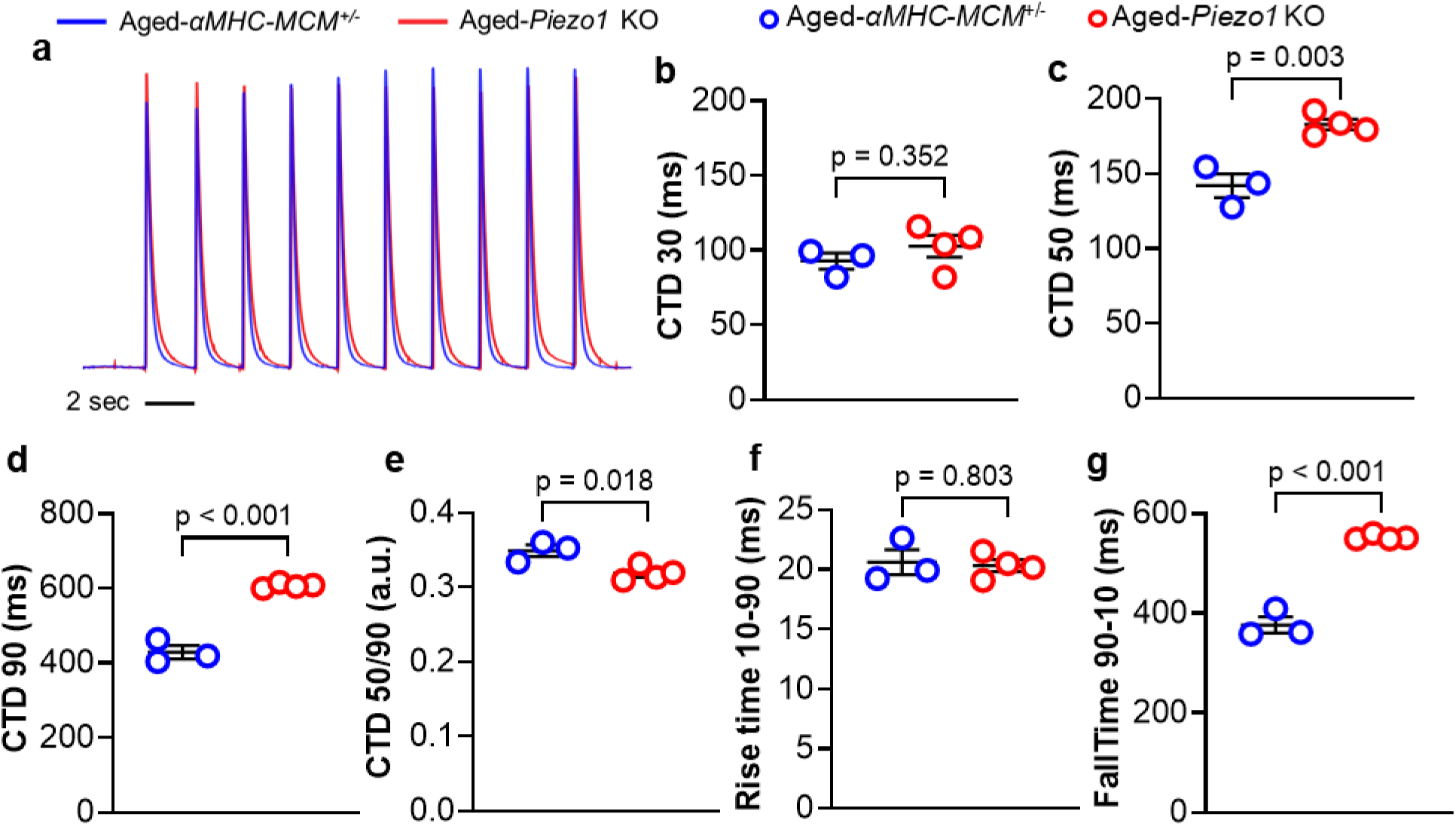
Altered Ca^2+^ handling in aged cardiomyocytes after deletion of *Piezo1*. (**a**) Raw traces illustrating Cacl-520 calcium transients in paced freshly isolated cardiomyocytes from aged *αMHC-MCM^+/-^*and *Piezo1 KO* mice, (**b**) calcium transient duration at 30% recovery (CTD30), (**c**) calcium transient duration at 50% recovery (CTD50), (**d**) calcium transient duration at 90% recovery (CTD90), (**e**) ratio of CTD 50 to CTD 90 (CTD 50/90), (**f**) rise time between 10% and 90% of peak transient (rise time 10-90), (**g**) fall time between 90% and 10% peak transient (fall time 90-10). Results are presented as mean ± SEM, Welch’s T-test, two-tailed was used between two groups.

### Altered Ca^2+^ handling in aged cardiomyocytes after deletion of Piezo1 is caused by dysregulation of Ca^2+^ handling proteins

We previously reported that PIEZO1 signalling influences several essential Ca^2+^ handling proteins in pressure-overload induced LVH^12^. Thus, we hypothesized that the altered Ca^2+^ handling kinetics observed in aged *Piezo1* KO cardiomyocytes could be driven by changes in critical Ca^2+^ handling proteins. No changes in mRNA expression of *SERCA2a, PLN, Slc8a1 or RyR2* were detected in aged *αMHC-MCM^+/-^*control hearts when compared to young *αMHC-MCM^+/-^* control hearts **(Fig. 6a-d).** However, when compared to aged *αMHC-MCM^+/-^*control hearts, aged *Piezo1* KO hearts showed significantly reduced mRNA expression of *SERCA2a and Slc8a1* (**Fig. 6a-b**). We found no differences in *PLN* or *RyR2* gene expression (**Fig. 6c-d**) between the aged groups.

**Figure. 6.**
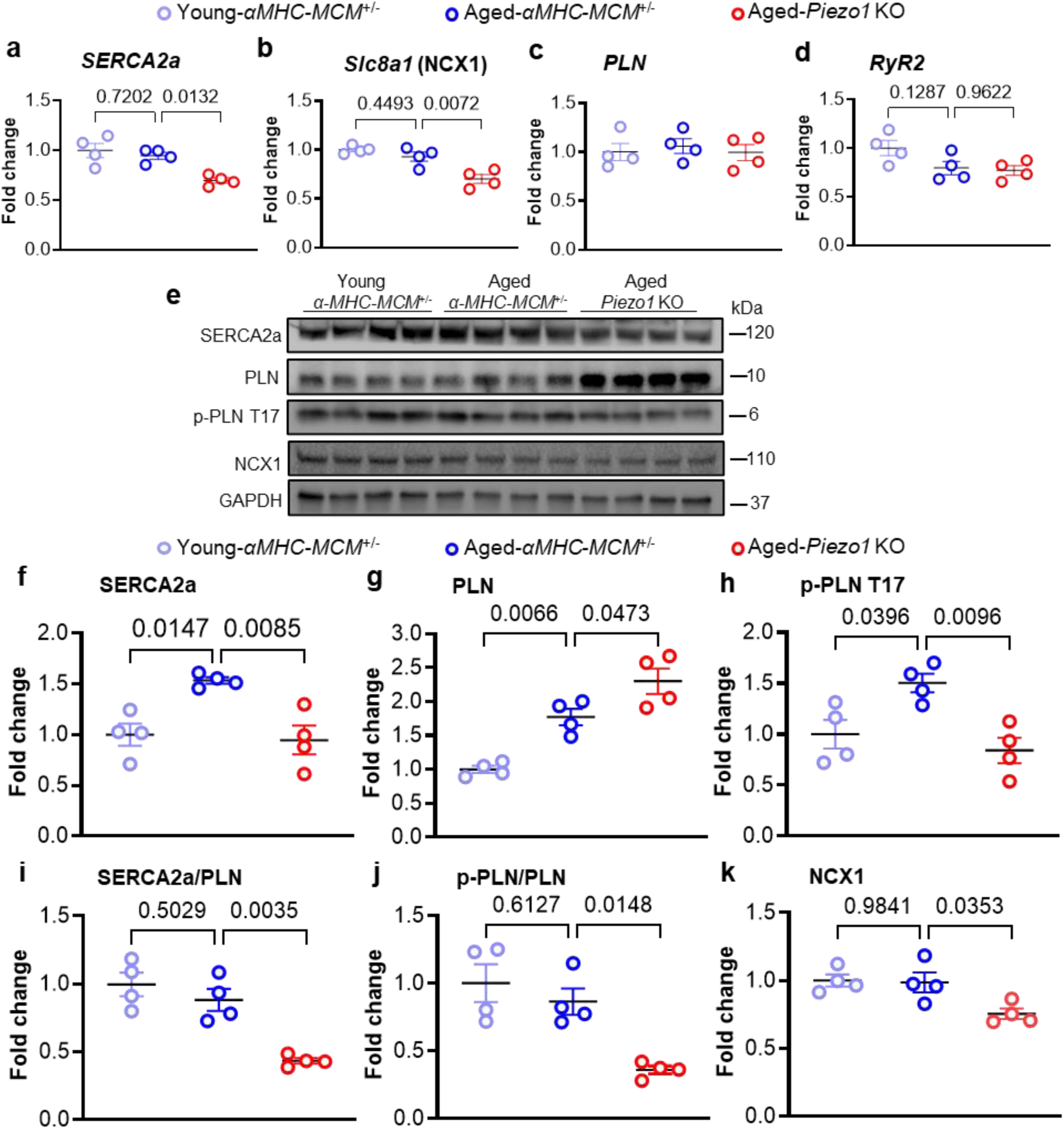
Altered Ca^2+^ handling in aged cardiomyocytes after deletion of *Piezo1* is caused by dysregulation of Ca^2+^ handling proteins. Selected Ca^2+^ gene or protein expression was measured in 3-month-old tamoxifen treated *αMHC-MCM^+/-^* mice (Young - *αMHC-MCM^+/-^*) and 21-month-old mice 18 months following tamoxifen treatment in both *αMHC-MCM^+/-^* mice (Aged - *αMHC-MCM^+/-^*) and *Piezo1* KO mice. (**a** - **d**) Relative mRNA expression of (**a**) *SERCA2a*, (**b**) *PLN*, (**c**) *Slc8a1*, (**d**) *RyR2*. The mRNA relative expression was normalized with *GAPDH* and calculated as fold change relative to 3-month-old tamoxifen treated *αMHC-MCM^+/-^* mice (Young-αMHC-MCM^+/-^ mice, n = 4 per group). (**e**) Representative western blots of SERCA2a, PLN, p-PLN T17 and NCX1 in LV tissue. (**f - k**) Quantitative data of (**f**) SERCA2a, (**g**) PLN, (**h**) p-PLN T17, (**i**) the ratio of SERCA2a/PLN, (**j**) the ratio of p-PLN T17/PLN and (**k**) NCX1were normalized with GAPDH. Fold changes were calculated relative to 3-month-old young-*αMHC-MCM^+/-^* mice (n = 4 per group). Results are presented as mean ± SEM, One-way ANOVA with Tukey’s post-hoc test for multiple comparisons was used.

We further assessed the expression of selected important Ca^2+^ handling proteins by western blot analysis (**Fig. 6e**). SERCA2a protein levels in aged *αMHC-MCM^+/-^* control hearts were 53% higher than in young *αMHC-MCM^+/-^* control hearts, suggesting increased SERCA2a protein expression with age (**Fig. 6f**). This upregulation of SERCA2a protein expression was not observed in the aged *Piezo1* KO hearts, which exhibited a 38% lower SERCA2a protein level when compared to age-matched *αMHC-MCM^+/-^* control hearts (**Fig. 6f**).

Phospholamban (PLN) expression in aged-*αMHC-MCM^+/-^* control hearts was 77% higher than in young *αMHC-MCM^+/-^* control hearts, while aged-*Piezo1* KO hearts showed a significant 30% increase in PLN protein levels when compared to aged-matched *αMHC-MCM^+/-^* control hearts (**Fig. 6g**). Phosphorylated PLN (Thr17) expression in aged-*αMHC-MCM^+/^*^-^ control hearts was 50% higher than in young *αMHC-MCM^+/-^* control hearts but this age-related increase was not observed in aged-*Piezo1* KO hearts (**Fig. 6h**). There were no significant differences in the SERCA2a to PLN ratio and the phosphorylated PLN (Thr17) to PLN ratio between young and aged *αMHC-MCM+/-* control hearts. In contrast, aged-*Piezo1* KO hearts exhibited significant reductions in both the SERCA2a to PLN ratio and the phosphorylated PLN (Thr17) to PLN ratio, by 51% and 58%, respectively, when compared to aged *αMHC-MCM^+/-^* control hearts (**Fig. 6i-j**). Additionally, the Na^+^/Ca^2+^ exchanger (NCX1) protein levels were 23% lower in aged-*Piezo1* KO hearts compared to aged-*αMHC-MCM^+/-^* control hearts (**Fig. 6k**). Together, these data indicate that the loss of *Piezo1* in adult cardiomyocytes results in significant dysregulation of key Ca^2+^ handling proteins as mice age.

### Cardiomyocyte-specific deletion of Piezo1 in adult mice results in age-induced bradycardia

While assessing cardiac function using echocardiography (**Fig. 4**), we observed a profound age-induced bradycardia in *Piezo1* KO mice (**Fig. 7a**). This reduced heart rate was the largest driver of the reduced cardiac output observed in *Piezo1* KO mice (**Fig. 4b**). The onset of the reduced heart rate in *Piezo1* KO mice was approximately six months after tamoxifen induction but was progressive with increasing age (**Fig. 7a**).

**Figure. 7.**
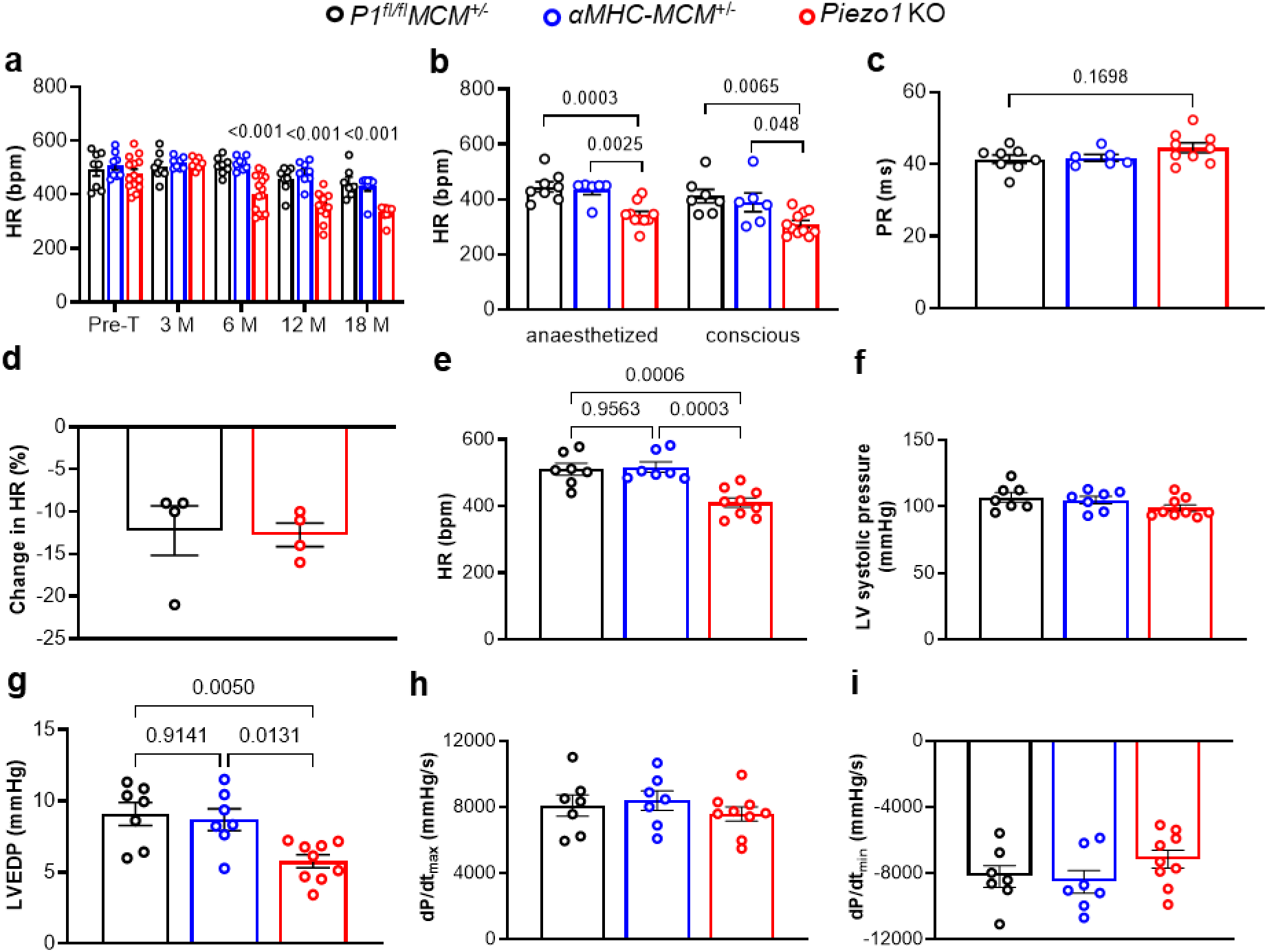
Deletion of *Piezo1* in cardiomyocytes results in significant bradycardia. (**a – d**) Echocardiographic measurements of (**a**) HR, heart rate from pre-treated (Pre-T) mice and 3 – 18 months following tamoxifen treatment in both *αMHC-MCM^+/-^* and *Piezo1* KO mice, and oil treated *P1^fl/fl^MCM^+/-^* controls (n = 7 – 14 per group). (**b**) Comparison of heart rate measured under anaesthesia using echocardiography or in conscious animals using the tail-cuff method in 21-month-old mice 18 months after oil treatment in *P1^f/l/fl^MCM^+/-^* mice or tamoxifen treatment in both *αMHC-MCM^+/-^* and *Piezo1* KO mice (n = 6 – 11 per group). (**c**) PR interval (n = 6 – 9 per group). (**d**) changes in HR measured after β-adrenoreceptor blockade with metoprolol (n = 4 per group). (**e - h**) Hemodynamic measurements in 21-month-old mice 18 months following tamoxifen treatment in both *αMHC-MCM^+/-^* and *Piezo1* KO mice: (**e**) HR, heart rate; (**f**) LV systolic pressure, left ventricular systolic pressure; (**g**) LVEDP, left ventricular end-diastolic pressure; (**h**) dP/dt_max_, the peak rate of left ventricular pressure increase; (**i**) dP/dt_min_, the peak rate of left ventricular pressure decrease (n = 7 – 9 per group). One-way or two-way ANOVA with Tukey’s post-hoc test for multiple comparisons was used to assess effects of genotype, age, and genotype by drug interaction. Results are presented as mean ± SEM.

To test whether the bradycardia observed in aged-*Piezo1* KO mice was influenced by anaesthesia, we also measured heart rates using the tail-cuff method in conscious mice and compared the results to those obtained under anaesthesia. Regardless of the method used, aged-*Piezo1* KO mice exhibited significantly lower heart rates than age-matched *P1^fl/fl^MCM^+/-^* and *αMHC-MCM^+/-^*control mice in both conscious and anesthetised conditions (**Fig. 7b**). Consequently, from the concomitant ECG recording the RR interval of aged *Piezo1* KO mice (182.4 ± 4.2 ms) was significantly longer when compared with age-matched *P1^fl/fl^MCM^+/-^* (154.3 ± 5.9 ms, p < 0.005) and *αMHC-MCM^+/-^* control mice (156.3 ± 7.1 ms, p < 0.02) but the PR interval was unchanged (**Fig. 7c**), indicative of sinus bradycardia.

Given the heart rate effects develop progressively over many months, it seemed unlikely these effects reflected a direct effect of the absence of PIEZO1 on the sinoatrial node pacemaker cells. This is consistent with previous short-term studies^12,13^ and the limited effect of the PIEZO1 agonist Yoda1 on isolated sinoatrial node tissue^10^. To ascertain whether the reduced heart rate observed in *Piezo1* KO mice could have been a result of alterations in sympathetic drive, heart rates were measured by echocardiography before and 20 minutes after the acute administration of a β-adrenergic receptor blocker (metoprolol, 1.0 mg/kg i.p.) in aged *P1^fl/fl^MCM^+/-^*control mice and *Piezo1* KO mice. The change in heart rate observed following β-adrenergic receptor blockade was comparable in both groups (**Fig. 7d**), suggesting that the lower baseline heart rate in the absence of PIEZO1 was not mediated by lower sympathetic drive.

Invasive hemodynamic measurements using left ventricular catheterization also confirmed that aged-*Piezo1* KO mice exhibited significant bradycardia when compared to either age-matched *P1^fl/fl^MCM^+/-^*or *αMHC-MCM^+/-^* control mice (**Fig. 7e**), without significant differences in left ventricular systolic pressure, dP/dt_max_ or dP/dt_min_ among the three groups (**Fig. 7f,h,i**). The left ventricular end-diastolic pressure (LVEDP) was significantly lower in aged-*Piezo1* KO mice when compared with age-matched *P1^fl/fl^MCM^+/-^* and *αMHC-MCM^+/-^*control mice (**Fig. 7g**).

### Aging-related bradycardia due to loss of Piezo1 in adult cardiomyocytes is linked to fibrotic chamber remodelling and decreased ANP levels

To investigate whether aging-induced bradycardia in *Piezo1* KO mice was associated with cardiac fibrotic remodeling, we used Masson’s trichrome staining to evaluate interstitial cardiac fibrosis in all four chambers of the heart (**Fig. 8a**). Aged *Piezo1* KO hearts and age-matched *αMHC-MCM^+/-^* control hearts were compared. Aged *Piezo1* KO hearts showed significantly higher levels of interstitial fibrosis (**Fig. 8b-e**). While the difference in the degree of fibrosis between groups was significantly higher in the left ventricle than the right ventricle, the most pronounced fibrotic phenotype was observed in the right atrium of the aged *Piezo1* KO hearts, with an average fibrotic area of 10.08% compared to 4.98% in age-matched *αMHC-MCM^+/-^* control hearts. In contrast, there was no difference between groups in the extent of fibrosis in the left atrium.

**Figure. 8.**
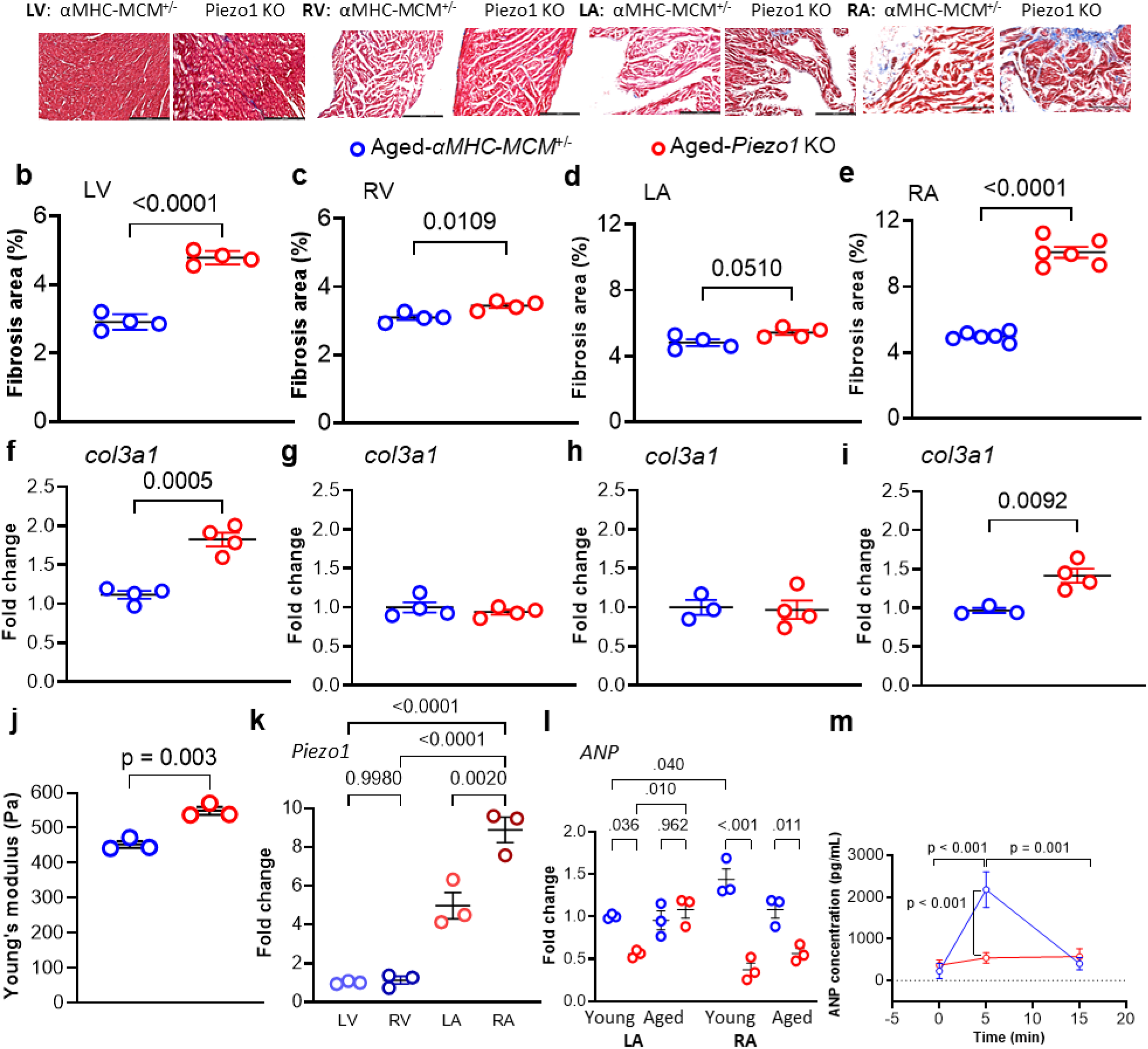
Aging induced bradycardia caused by loss of PIEZO1 in adult cardiomyocytes is associated with chamber remodelling and reduced ANP levels. (**a**) Representative micrographs of Masson’s trichrome staining of left and right ventricular and left and right atrial sections from 21-month-old mice, 18 months following tamoxifen treatment in both *αMHC-MCM^+/-^* and *Piezo1* KO mice, scale bar, 200 µm. (*b* - *e*) Quantitation of Masson’s trichrome staining of (*b*) LV, (*c*) RV, (*d*) LA, (*e*) RA (n = 4 - 6 per group). (*f* – *i*) Relative collagen III (*Col3a1*) mRNA expression of (*f*) LV, (*g*) RV, (*h*) LA, (*i*) RA (n = 3 per group). The mRNA relative expression was normalized with *GAPDH* and calculated as fold change to *αMHC-MCM^+/-^* mice. *(j)* Cardiomyocyte stiffness measurements using atomic force microscopy (n = 3 per group). Results are presented as mean ± SEM, Welch’s T-test, two-tailed was used between two groups. (*k*) Relative Piezo1 mRNA expression in LV, RV, LA and RA isolated cardiomyocytes from 3-month-old *αMHC-MCM^+/-^* mice (n = 3 per group). The mRNA relative expression was normalized with *GAPDH* and calculated as fold change relative to the values of LV cardiomyocytes. (**l**) Relative *ANP* mRNA expression in LA and RA tissues from 3-month-old mice following tamoxifen treatment in both *αMHC-MCM^+/-^*mice and *Piezo1* KO mice, and 21-month-old mice 18 months following tamoxifen treatment in both *αMHC-MCM^+/-^* and *Piezo1* KO mice (n = 3 per group). The mRNA relative expression was normalized with *GAPDH* and calculated as fold change relative to the values of LA tissue of 3-month-old tamoxifen treated αMHC-MCM^+/-^ mice (Young). (**m**) Plasma ANP levels were measured at 0, 5, and 15 min after acute expansion of intravascular volume in 3-month-old tamoxifen treated *αMHC-MCM^+/-^* and *Piezo1* KO mice (n = 7 per group).

Analysis of the mRNA expression level of Collagen III (*col3a1*), a major marker of myocardial fibrosis, in each heart chamber revealed a significant increase in the left ventricle (63%) and right atrium (46%) of aged-*Piezo1* KO hearts when compared with age-matched *αMHC-MCM^+/-^*control hearts (**Fig. 8f-i**). To further characterise the fibrotic remodelling, we also assessed the mechanical properties of the ventricular myocytes using atomic force microscopy. Aged *Piezo1* KO myocytes were found to be significantly stiffer than age-matched *αMHC-MCM^+/-^* myocytes (**Fig. 8j**).

Given the chamber-specific effects of the observed fibrotic remodelling, we asked the question of whether *Piezo1* levels may also differ throughout the myocardium. We performed reverse transcription quantitative PCR (RT-qPCR) on freshly isolated cardiomyocytes from young *αMHC-MCM^+/-^* hearts. The highest *Piezo1* levels were observed in the right atrium, approximately 8 times greater than in the left and right ventricles and 1.8 times greater than in the left atrium (**Fig. 8k**).

It is well documented that atrial natriuretic peptide (ANP) has anti-fibrotic functions in the heart^32–34^. Given ANP is released by a still unknown stretch-dependent pathway^35^, we first asked the question of whether deletion of cardiomyocyte *Piezo1* influenced *ANP* levels. We measured *ANP* mRNA expression levels in left and right atrial tissue from both young and aged-*αMHC-MCM^+/-^* controls and *Piezo1* KO hearts. We found that the loss of *Piezo1* resulted in a 44% reduction in *ANP* expression in the left atrium of young-*Piezo1* KO hearts (p < 0.05), but *ANP* expression levels returned to control levels in aged-*Piezo1* KO left atria. In young-*αMHC-MCM^+/-^* hearts, *ANP* mRNA expression was significantly higher in the right atrium than in the LA. However, young-*Piezo1* KO hearts exhibited decreased *ANP* mRNA expression in the right atrium, and this reduction persisted with age (**Fig. 8l**). These findings indicate that deletion of cardiomyocyte *Piezo1* influences *ANP* levels.

To further investigate whether the deletion of cardiomyocyte *Piezo1* impairs ANP release *in vivo*, we performed an acute intravascular volume expansion in young adult *αMHC-MCM^+/-^* control mice and *Piezo1* KO mice. In *αMHC-MCM^+/-^* mice, a 60% increase in total circulating blood volume triggered a substantial elevation in plasma ANP, rising from a mean value of 227 ± 184 pg/mL to 2176 ± 428 pg/mL within 5 minutes. This level subsequently decreased to 405 ± 148 pg/mL by 15 minutes. In contrast, the increase in ANP in response to volume expansion was blunted significantly in *Piezo1* KO mice. At 5 minutes post-infusion, the mean ANP level in *Piezo1* KO mice was 544 ± 127 pg/mL, only 25% of the peak value in *αMHC-MCM^+/-^* control mice (**Fig. 8m**).

Together, these findings suggest that the deletion of cardiomyocyte *Piezo1* in adult hearts leads to significant age-induced right atrial fibrotic remodelling and reduced levels of the anti-fibrotic protein, ANP. Moreover, the *in vivo* volume loading data are consistent with involvement of PIEZO1 in the stretch-induced release of ANP.

## Discussion

In this work we looked at the long-term impact of *Piezo1* deletion in adult cardiomyocytes using an inducible *Piezo1* KO mouse model. Previous work had demonstrated that the loss of cardiomyocyte *Piezo1* early on in development using an *MLC2v-Cre* driver resulted in a dilated left ventricle and impaired cardiac function^11^. In our study, using serial echocardiography, deleting *Piezo1* in adult cardiomyocytes using a tamoxifen inducible *Myh6-Cre* driver did not result in a dilated cardiomyopathy in *Piezo1* KO hearts even 18 months after *Piezo1* deletion. Thus, it would seem likely that the phenotype observed by Jiang *et al.* using an *MLC2v-Cre* stems from influences on early cardiac development that provide a template for the dilated phenotype. The influence of the loss of cardiomyocyte PIEZO1 on cardiac remodelling in this study and others^11,13^ strongly suggests loss of function *Piezo1* variants may have deleterious cardiac effects magnified by aging.

The first obvious effect we noted was increased mortality of *Piezo1* KO mice during aging. This effect was profound and resulted in earlier cessation of the study, which was originally planned to extend to 24 months post *Piezo1* deletion. This early mortality coincided with increased expression of the senescence associated secretory phenotype (SASP) proteins, including *IL1β, IL6, TGFβ, and TNFα*, indicative of accelerated senescence, which contributes to the aging phenotype^25,36,37^. PIEZO1 has previously been linked to senescence in skeletal muscle stem cells through P53 signalling^38^, and we also observed PIEZO1-dependent effects on P53 as well as P16 and P21. Previous work in skeletal muscle links this to signalling through reactive oxygen species (ROS) and indeed initial work in cardiomyocytes suggests PIEZO1 can influence ROS-signalling in cardiomyocytes^11^.

During aging in both mice and human subjects, there is a notable hypertrophic response in the myocardium previously linked to mechanistic target of rapamycin (mTOR) and insulin-like growth factor-1 (IGF-1) signalling^39,40^. We indeed observed a hypertrophic response during aging. However, when we compared aged-*Piezo1* KO mice hearts to their age-matched controls, we noticed significant blunting of the age-induced hypertrophy. We showed by isolating individual myocytes that they were smaller in aged-*Piezo1* KO mice compared to controls. We had previously shown that PIEZO1 drives a CaMKII-dependent signalling pathway in response to pressure-overload to instigate left ventricular hypertrophy^12^. In aged-*Piezo1* KO hearts, we found evidence of reduced CaMKII activation consistent with the reduction in age-related hypertrophy we observed in *Piezo1* KO hearts. This implicates CaMKII activation in aging induced hypertrophy. Previous work in many organisms suggests CaMKII can contribute to cardiac remodelling during aging^41,42^. We surmise, therefore, that PIEZO1 is acting as a homeostatic regulator during aging. Its loss reduces cardiac mass by reducing the activation of CaMKII. In fact, our observation in cardiac aging has parallels with the role of PIEZO1 in bone, where it has been shown to oppose age-associated cortical bone loss^43^.

When we examined the cardiac functional effects of *Piezo1* deletion, we found little impact on left ventricular systolic function with aging but a profound effect on diastolic left ventricular function due to impaired diastolic relaxation, and this was consistent with the impact of *Piezo1* deletion on cardiomyocyte Ca^2+^ handling kinetics and key Ca^2+^ handling proteins.

We also observed a profound age-induced bradycardia in *Piezo1* KO mice. If the bradycardia were due to an inherent role of PIEZO1 in cardiac pacemaker cells, we would expect to observe this response soon after *Piezo1* deletion. Instead, we observed a progressive reduction in heart rate with aging over many months. This observation suggests that the observed bradycardia is a secondary effect of the loss of PIEZO1 that develops progressively with aging. Consistent with this hypothesis, previous work suggests that while the sinoatrial node may respond to mechanical cues^44^, this seems to be independent of PIEZO1 because addition of the small molecule PIEZO1 agonist Yoda1 has little to no effect on sinoatrial node function^10^. The observed age-induced bradycardia in *Piezo1* KO mice was also not driven by reduced sympathetic drive because treatment of aged mice with a β-adrenergic receptor blocker resulted in the same relative change in heart rate regardless of the genotype.

The most likely candidate to explain the progressive bradycardia with aging in the absence of PIEZO1 would appear to be our parallel observation of increased cardiac fibrosis, which was most prominent in the right atrium of aging *Piezo1* KO mice. Fibrotic remodelling within the sinoatrial node has previously been linked to the development of the tachycardia-bradycardia syndrome and its negative clinical sequelae^45^. The fibrotic remodelling we identified could be related to the reduced anti-fibrotic action of atrial natriuretic peptide (ANP)^33,34,46^. We identified a profound reduction in *ANP* mRNA levels in *Piezo1* KO mice, and this was particularly notable in the right atrium relative to the left atrium with aging.

ANP is released from cardiomyocytes in response to atrial stretch, although the exact mechanism by which atrial stretch induces ANP release has remained a mystery^35,47^. Using an *in vivo* volume expansion assay, we showed that *Piezo1* KO mice not only have lower mRNA levels of ANP but that the atrial stretch-induced release of ANP is significantly blunted in the absence of PIEZO1. This has profound implications for our understating of stretch-induced regulation of blood volume through ANP in normal physiology. This also provides a mechanism whereby long-term loss of PIEZO1 could result in remodelling within the atria, through a reduction in the anti-fibrotic action of ANP, thus influencing sinoatrial node function leading to bradycardia.

In conclusion, we show that cardiomyocyte PIEZO1 plays a critical homeostatic role in cardiac aging. The loss of PIEZO1 blunts the normal cardiac hypertrophic response to aging, adversely impacts diastolic cardiac function, reduces ANP levels and causes progressive cardiac fibrosis with bradycardia, and shortens life expectancy. These findings provide the first evidence that PIEZO1 plays a key role in cardiac aging, and the detailed molecular mechanisms of its protective actions may reveal novel anti-aging strategies.

## Acknowledgements

The authors gratefully acknowledge funding from the National Health and Medical Research Council (NHMRC) of Australia to BM and MF through a project grant (APP2010464) and an NSW Cardiovascular Disease Senior Scientist Grant to BM. CC is supported by an ARC Future Fellowship.

## Author contributions

ZYY, Data curation, Formal analysis, Validation, Investigation, Methodology, Writing – original draft, Project administration; YG, Conceptualization, Data curation, Formal analysis, Investigation, Visualization, Writing - original draft; SK, Data curation; DC, HL, JW, EN, Data curation, Methodology; PM, MX, Resources, Writing – original draft; CC, MF, BM, Conceptualization, Resources, Supervision, Funding acquisition, Validation, Methodology, Writing - original draft, Project administration.

